# Genetic Variation as a Long-Distance Modulator of *RAD21* Expression in Humans

**DOI:** 10.1101/2021.05.27.445915

**Authors:** William Schierding, Julia A. Horsfield, Justin O’Sullivan

## Abstract

Mutations and changes in expression in *RAD21* are common across cancers types and outside of cancer can result in cohesinopathy. As such, exploration of variants that modify *RAD21* enhancer activity, across the genome, may also provide insights into mechanisms by which distinct variants impact healthy human development and disease. We searched 42,953,834 genomic variants for a spatial-eQTL association with the transcription of *RAD21*. We identified 123 significant associations (FDR < 0.05), which are local (*cis*) or long-distance (*trans*) regulators of *RAD21* expression. The 123 variants co-regulate a further seven genes, enriched for having Sp2 transcription factor binding sites in their promoter regions. The Sp2 transcription factor and six of the seven genes had previously been associated with cancer onset, progression, and metastasis. Our results suggest that genome-wide variation in non-coding regions impacts on *RAD21* transcript levels in addition to other genes, which then could impact on oncogenesis and the process of ubiquitination. This identification of distant co-regulation of oncogenes represents a strategy for discovery of novel genetic regions which impact cancer onset and a potential for diagnostics.

## 1. Introduction

Spatial organization and compaction of chromosomes in the nucleus forms a higher order chromatin structure that involves non-random folding of DNA on different scales, including chromosome compartments, topologically associated domains (TADs), and loops. Cohesin participates in genome organisation by forming looped domains with the CCCTC binding factor (CTCF) via CTCF’s ability to serve as loop anchors and block loop extrusion by cohesin^1^. The human cohesin complex contains four integral subunits: two structural maintenance proteins (SMC1A, SMC3), one stromalin HEAT-repeat domain subunit (STAG1 or STAG2), and one kleisin subunit (RAD21)^2^. Alteration of the components of cohesin alter high order chromatin structure and thus affect various biological processes within the nucleus. Thus, a complete loss of cohesin is not tolerated because of its essential role in cell division during normal tissue development^3^. For example, *RAD21* and the other cohesin genes are mutationally constrained, with a high probability of loss-of-function intolerance (pLI > 0.9; LOEUF < 0.35)^4^. In contrast, mutations that lead to reduced cohesin dose are survivable, but can alter gene expression^5^ leading to developmental disorders (known as ‘cohesinopathies’)^6^ and cancer^7–13^.

Remarkably, features of the predominant cohesinopathy, Cornelia de Lange Syndrome (CdLS), can exhibit with less than a 30% depletion in NIPBL protein levels (NIPBL loads cohesin onto DNA)^14^. Similarly, mutations in *RAD21* leading to a cohesinopathy typically affect the *RAD21* interface with the other cohesin proteins^15^. In cancer, alteration of *RAD21* expression is closely related to the development, prognosis, invasion, and metastasis of cancer cells^16–18^. Additionally, non-coding mutations around the *RAD21* locus have been found by genome wide association studies (GWAS) to track with multiple disease phenotypes^19^. However, the GTEx catalogue lists thousands of genetic variants in healthy individuals that are associated with local control of *RAD21* expression (*cis*-eQTLs)^20^. Our previous work found that many of these variants in *cis*-eQTLs are also associated with co-regulation of a network of genes involved in complex disease processes^21^. Our results indicated spatial eQTLs as identifying disease associations through mechanisms alternative to those disease associations found via GWAS. While GWAS reports susceptibility loci for a phenotype, traditional GWAS has several limitations, including an inability to discern a causal variant within the many linked SNPs in the risk locus. This may prevent true gene–trait associations from being identified. By integrating regulatory mechanisms such as spatial associations and eQTLs, which measure the ability of a variant to regulate the expression of target genes, the confluence of these data types can illuminate the underlying influence on the process of pathogenesis. Thus, eQTLs dissect genetic mechanism of variants implicated in multiple diseases and to prioritize SNPs or genes for further functional experiments.

We hypothesised that by again leveraging these regulatory associations, we could elucidate cohesin-associated pathologies within control elements across the genome which are affected by subtle, combinatorial changes in the regulation of *RAD21*. Here, we link the three dimensional (3D) structure of the genome with eQTL data to identify common variants within the genome that affect the transcriptional levels of *RAD21*. Some of these eQTLs have previously been associated with disease pathways. However, their regulatory relationships with *RAD21* open up novel avenues for exploration of disease onset for both cohesinopathy disorders and cancer.

## 2. Results

### Variant Selection for Genome-wide search for distant regulators of transcription

To test whether variants across the genome had a significant effect on the transcription of *RAD21* we performed a genome-wide search of all 42,953,834 SNPs in dbSNP151 (as available in GTEx v8^20^) for a spatial-eQTL association with the *RAD21* transcript level (Figure 1; Table 1). We identified 123 SNPs that were significantly associated with *RAD21* transcript levels from various genomic distances (120 *cis*, 1 *trans*-intrachromosomal, and 2 *trans*-interchromosomal; Figure 2a; FDR < 0.05). Of note, the *trans*-intrachromosomal connection was with a locus that was > 16 Mb away from *RAD21* within the linear sequence of chromosome 8 in eQTL tissue from the adrenal gland. The *trans*-interchromosomal connections involved SNPs that are located on chromosomes 11 and 13, in eQTL tissues from the hypothalamus (chr 11) and transformed lymphocytes (chr 13).

**Figure 1.**
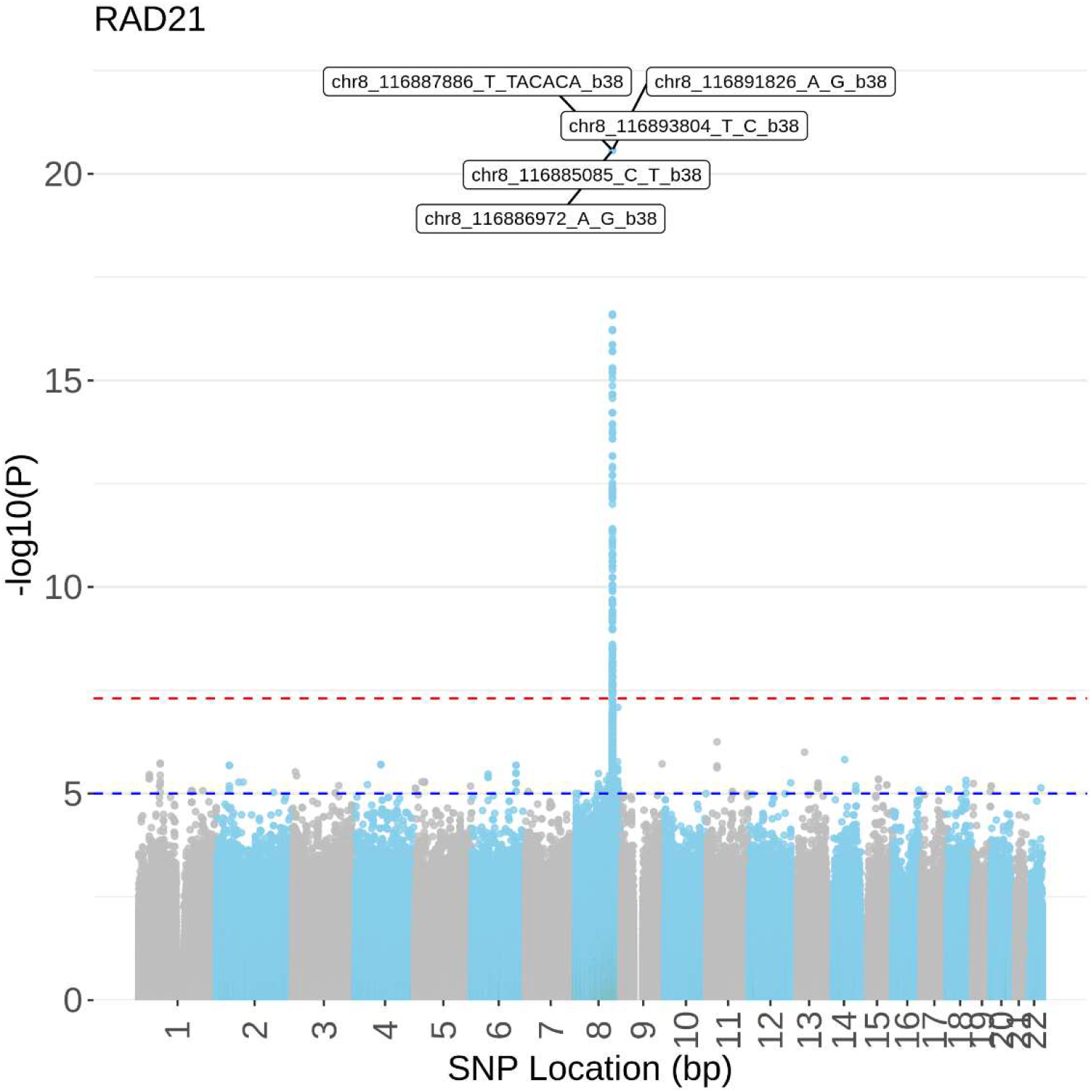
Manhattan plot of all eQTL p-values between genome-wide SNPs and RAD21 expression across all GTEx tissues. The top 5 SNPs are highlighted. Despite there being no similar standard for genome-wide eQTL studies, the red (p < 5 × 10^−8^) and blue (p < 1 × 10^−5^) dashed lines represent the typical p-value thresholds from GWAS studies as a reference.

**Figure 2.**
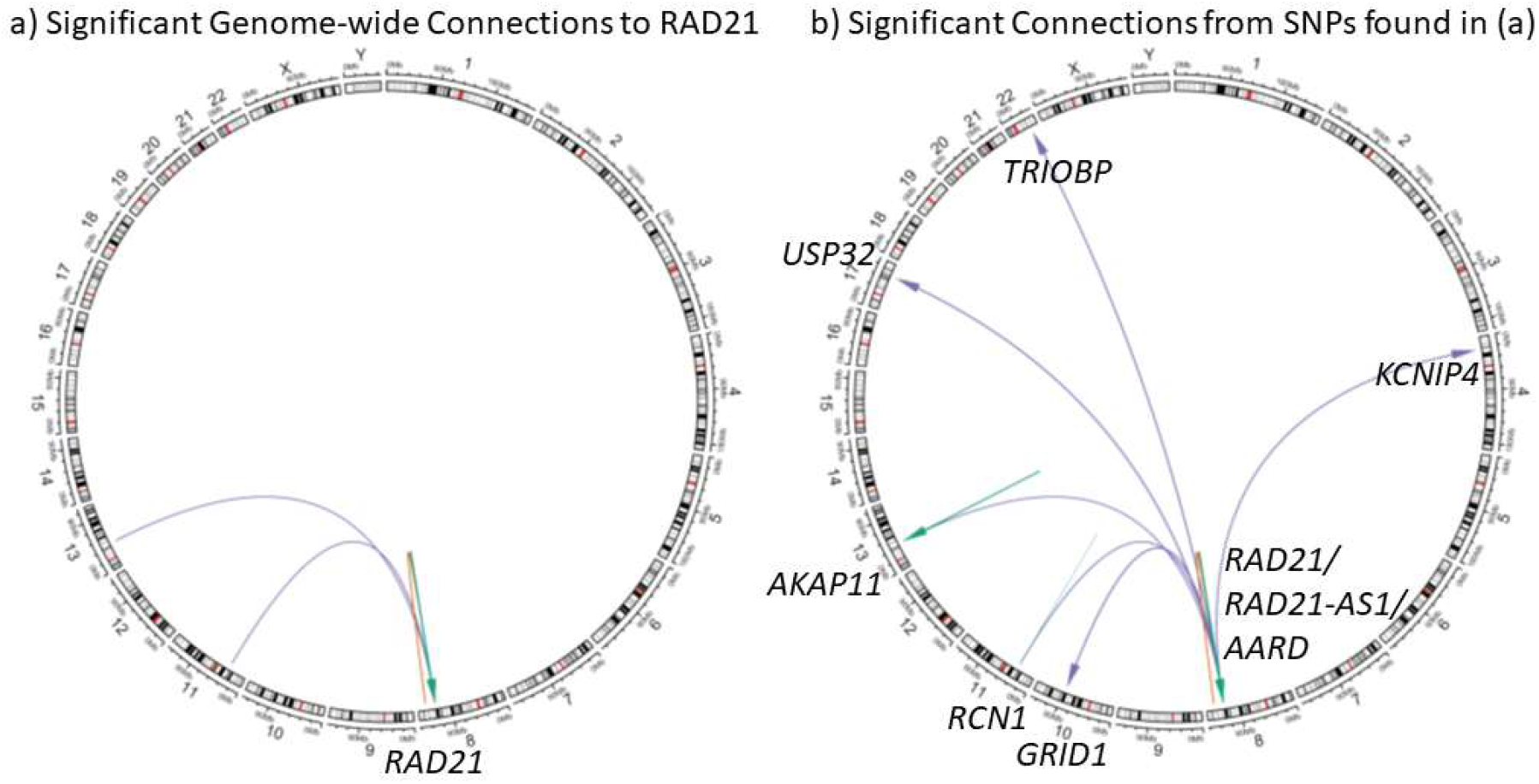
Circos plots showing the interactions and locations of the genes found for: a) Significant Genome-wide Connections to RAD21. b) Significant Connections from SNPs found in (a). This resulted in the discovery of a hub of eight co-regulated genes: AARD, *AKAP11, GRID1, KCNIP4, RCN1, TRIOBP*, and *USP32*. Color of the lines represents the distance of the connection, where green = *cis*, orange = *trans*-intrachromosomal, and purple = *trans*-interchromosomal. Arrows represent direction of regulation (SNP to gene).

### Identification of regulatory potential of distal SNPs with *RAD21* Spatial Relationships

To test these 123 SNPs for their potential to co-regulate other genes both nearby and distal, all SNPs with significant SNP-*RAD21* spatial-eQTL relationships (FDR < 0.05) were then tested with the CoDeS3D algorithm^22^ to discover other genes that were potentially co-regulated by these SNPs (Figure 2b; Table 2). This resulted in the discovery of a hub of eight co-regulated genes: AARD, *AKAP11, GRID1, KCNIP4, RAD21, RCN1, TRIOBP*, and *USP32* (Figure 2b). The two SNPs (rs238256 and rs10639528) with *trans*-interchromosomal connections to *RAD21* are responsible for the associations with *AKAP11* and *RCN1* transcript levels. Notably, SNPs regulating *RAD21* in *cis* were associated with a *trans*-interchromosomal eQTL connection with *GRID1*. Of note, *GRID1* is <500 kb away from *WAPL* on chromosome 10q23.2 (WAPL protein facilitates cohesin’s removal from chromatin).

### Gene set enrichment analysis implicates the Sp2 transcription factor as a common regulator on these gene promoters

The promoter regions (+/−1kb from TSS) of *RAD21* and the seven co-regulated genes we identified in this analysis were all significantly enriched for Sp2 regulatory motifs (Table 3; GGSNNGGGGGCGGGGCCNGNGS; Transfac putative transcription factor binding site M09658; p=.03047). This suggests that the level at which Sp2 acts is upstream of *RAD21* and the co-regulated genes. Sp2 is a DNA binding transcription factor in the Sp subfamily, which are required for the expression of cell cycle- and developmentally-regulated genes, and deregulated expression Sp family members is associated with human tumorigenesis^23^. Notably, it has recently been found that Sp2 is significantly upregulated in cohesin (*STAG2*) mutant CMK cells following 4 hours of WNT3A treatment^24^.

### AKAP11 and USP32 are Linked to Regulation of Cohesin via STRING Protein-Protein Interaction Networks

STRING analysis (Figure 3) identified that two genes in this study have been shown to be coexpressed (*KCNIP4* and *GRID1;* RNA co-expression score = 0.135)^25^. Additionally, *AKAP11* and *RAD21* are linked through a common co-expression with cohesin genes *PDS5A* (RNA co-expression score = 0.208) and PDS5B (RNA co-expression score = 0.203). Finally, *RAD21* and *USP32* are linked through a high-confidence Protein-Protein interaction between USP32 and SMC1A^26^ (SMC1A is part of the cohesin complex with RAD21).

**Figure 3.**
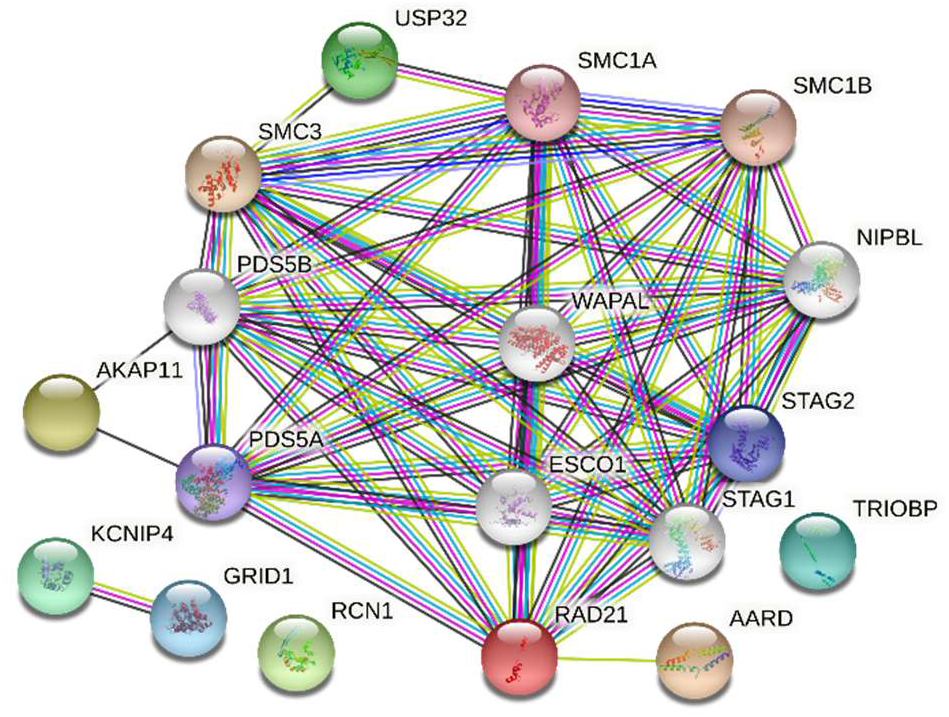
STRING analysis identified a connected network linking USP32, AKAP11, and AARD to the cohesin proteins including RAD21. Additionally, two genes in this study have been shown to be coexpressed (*KCNIP4* and *GRID1;* RNA co-expression score = 0.135).

### The Cohesin Complex and its Co-Regulated Genes are Enriched for Loss-of-Function Intolerance

The gnomAD catalog categorizes the probability of loss-of-function (LoF) mutation intolerance (pLI) as a pLI ≥ 0.9 with 3,063 out of 19,704 genes (15.5%) having LoF-intolerance^4^. gnomAD also reports the observed/expected score 90% confidence interval for continuity across the spectrum of selection that can distinguish selection from low sample size bias. Using the 90% upper bound of the loss-of-function confidence interval (LOEUF), LoF mutation intolerance is defined as LOEUF < 0.35. For example, the LOEUF scores can differentiate the 678 genes essential for human cell viability (mean LOEUF of 0.63) compared to the 777 non-essential genes (mean LOEUF of 1.34). In gnomAD, 6282 genes (31.8%) have a LOEUF < 0.63 and 4897 genes (24.9%) have a LOEUF > 1.34.

It is well established that the cohesin complex is necessary for healthy development. Consistent with this, LoF mutations within the *RAD21, SMC1A, SMC3, STAG1, STAG2, CTCF, MAU2, NIPBL, PDS5A, PDS5B*, and *WAPL/WAPAL* genes occur far less than expected, as indicated by the pLi > 0.99 and LOEUF < 0.256 for all 10 cohesin component genes (Table 4). Of the seven co-regulated genes we identified in our analysis, three (42.9%) were LoF intolerant (Table 4; AKAP11 pLI=0.97878; GRID1 pLI=0.99931; and USP32 pLI=1.0). The other four were somewhat tolerant (AARD: pLI=0.32580; KCNIP4 pLI=0.82064) or completely LoF-tolerant (RCN1: pLI=2.8821×10^−5^; TRIOBP: pLI=2.0161×10^−28^). Including *RAD21*, that is 4 out of 8 genes in the analysis with LoF-intolerance, a significantly larger proportion than expected by chance based on pLI (p=0.02, fisher exact test). Notably, the *SP2* gene is also LoF-intolerant (pLI=1.0). Using LOEUF, 4 of the 7 co-regulated genes are below the average essential gene LOEUF score (0.63) while none are above the average non-essential gene LOEUF (1.34) (Table 4). SP2 is also below 0.63.

## 3. Discussion

An insufficiency of *RAD21* that is not lethal can contribute to cohesinopathy or cancer. Here, we searched for genomic variation with subclinical moderation of *RAD21* transcript levels to explore how modification of *RAD21* enhancers across the genome could impact healthy human development and disease. Amongst the 42,953,834 genomic variants that mark the genome, we found significant (FDR < 0.05) spatial-eQTL associations with transcription of *RAD21* in 123 variants. This analysis identified many local (*cis*) regulatory regions, but also three loci that were long-distance regulators of *RAD21* expression. Overall, these 123 variants co-regulated a further seven genes, whose promoters were enriched for Sp2 transcription factor binding sites. These results highlight the potential roles of transcription factor Sp2 and deubiquitination (via *USP32*) in *RAD21* transcript level regulation.

### The Role of Sp2 as a Common Transcription Factor

Sp2 primarily localizes at CCAAT motifs, and the CCAAT box binding transcription factor Nf-y is the major partner for Sp2-chromatin interactions^27^. The Sp2 transcription factor gene ontology (GO) annotations include DNA-binding transcription factor activity, RNA polymerase II-specific, and histone deacetylase binding. Sp2 is almost universally expressed across all human tissues. Although the role of Sp2 in normal human tissues has not been well examined, Sp2 knockouts in zebrafish are embryonically lethal^23^ indicating that it is an essential gene, which is also supported in humans via the high pLI in gnomAD (pLI = 0.99721). We propose that the enrichment of these seven genes with Sp2 transcription factor binding sites results from one of two scenarios:

1. modification of *RAD21* expression alters Sp2 regulation, the downstream effects of which modify the transcript levels of these co-regulated genes.
2. modification of Sp2 itself alters pathways upstream of the seven co-regulated genes, of which *RAD21* is one.

As it has recently been shown that the knockdown of STAG2 in CMK cells was correlated with an upregulation of Sp2, it would appear that (1) would favoured, as this would argue a direct correlation between cohesin levels and Sp2^24^. However, as *RAD21* and its co-regulated genes all have Sp2 binding sites (and are therefore potentially co-regulated by Sp2), there is also evidence that Sp2 is the intermediary between an undetermined upstream signal, *RAD21*, and the co-regulated genes.

### The Role of Deubiquitination and USP32

In our analysis, three SNPs with *cis*-regulatory relationships with *RAD21* also co-regulated *USP32*, a ubiquitin-specific protease. Protein ubiquitylation is a post-translational modification with an important role in regulating protein degradation. Ubiquitylation is a reversible process, removed by deubiquitylating enzymes, of which one class is the ubiquitin-specific proteases (USPs). Ubiquitination of the cohesin complex helps to regulate chromosome segregation and cohesion during mitotic progression through control of replication fork integrity and the cellular response to replication stress^28,29^. While previous findings have pointed to Ubiquitin-specific proteases USP13 and USP37, the mechanisms and effects of cohesin ubiquitination remain largely undefined^29^. Of note, in mitosis RAD21 is poly-ubiquitinated (targeting RAD21 for degradation), but the role of USPs in this hasn’t been fully elucidated. Thus, USP32, a poorly characterized deubiquitinating enzyme, might also play a role. For example, both USP37 and USP32 can reduce resistance to cisplatin-targeting therapies in cancer^30^, suggesting the involvement of USP32 in drug resistance. Further evidence to this role of USP32 and cohesin has been found in a screen of deubiquitinating enzyme interactions, where USP32 had a high-confidence Protein-Protein interaction with SMC1A^26^ (which has been found to be mono-ubiquitinated in mitosis^29^). Additionally, *USP32* mutations have been found in human leukemia cells, which frequently carry recurrent cohesin deficits, including mutations in SMC1^31^ and RAD21^32^.

The role of USP32 in cohesin regulation could also be implicated through mutual calcium-dependence. While not significant in the gene set enrichment, it is notable that *KCNIP4, RCN1*, and *USP32* are all involved in the same molecular function: calcium ion binding. The role of calcium with USP32 is undefined, but USP6 (97% sequence identity with USP32 as a chimeric fusion of USP32 and TBC1D3^33^) induces tumorigenesis in a calcium-dependent manner through interaction with the ubiquitous calcium-binding protein calmodulin^34^. This calcium-ion dependence could be important in the co-regulation of *RAD21* activity, as one mechanism for the dissolution of the cohesin complex from chromatids is through calpain-1, a RAD21 peptidase that cleaves RAD21 at L192 in a calciumdependent manner^35^.

### Role of RAD21-associated genes in Cancer

In total, six of the seven co-regulated genes and Sp2 have connections to cancer onset, progression, and metastasis. Overall, results suggest that genome-wide variation in enhancer regions could impact on cohesin function as well as other co-regulated genes which could impact on oncogenesis.

The enrichment for LoF-intolerance amongst the *RAD21* co-regulated genes could be associated with an importance of these genes to normal development and cancer. The three LoF-intolerant genes (*AKAP11, GRID1*, and *USP32*) are all mutated in 5-8% of all Endometrial cancers in TCGA^36^. GRID1 is also at the breakpoint of several structural translocations such as t(10;20)(q23;q13) DPM1/GRID1 in Acute Myeloid Leukemia^37^. USP32 is overexpressed in breast cancer and human small cell lung cancer and may serve as an oncogene through promoting cell proliferation and tumor metastasis^38,39^. Many AKAP proteins, including AKAP11, are also commonly mutated in breast cancer^40^.

Beyond the three LoF-intolerant genes, three other *RAD21* co-regulated genes also have putative roles in cancer. TRIOBP has been identified in a range of different cancers including lung carcinoma, glioblastoma, esophageal, pancreatic, prostate, lung, and breast cancer^41^. RCN1 inhibits IP3R1-mediated ER calcium release and thereby inhibiting ER stress-induced apoptosis, which is disrupted by mutation in breast, colorectal, kidney, and liver cancer^42^. The KCNIP4 gene is disrupted by a translocation (t(3;4)(p13;p15)) in renal cell cancer^43^ and by a mutation in squamous cell lung cancer^44^.

In cancer, the expression pattern of Sp2 is inversely correlated with CEACAM1 expression in prostate cancer cells, likely due to its role in recruiting histone deacetylase to the CEACAM1 promoter (downregulation of a tumor suppressor gene)^45^. More recently, Sp2 disruption was shown to promote invasion and metastasis of hepatocellular carcinoma, possibly through TRIB3^46^.

### Conclusion

By exploring 42,953,834 variants across the genome, we identified 123 that modify *RAD21* transcript levels, and also co-regulate 7 genes. These genes share key molecular pathways with RAD21, including being regulated by transcription factor Sp2 binding, having a role in deubiquitination, and oncogenesis. In addition, *RAD21* and its co-regulated genes are enriched for being mutationally constrained. This work demonstrates that co-regulation of *RAD21* with associated genes is likely to be functionally important, and has potential for disruption in pathologies such as cancer.

## 4. Materials and Methods

### Variant Selection for Genome-wide search for distant regulators of transcription

To test whether variants across the genome had a significant effect on the transcription of *RAD21* we performed a genome-wide search of all 42,953,834 SNPs in dbSNP151 (as available in GTEx v8^20^) for an association with transcription of *RAD21* (Figure 4).

**Figure 4.**
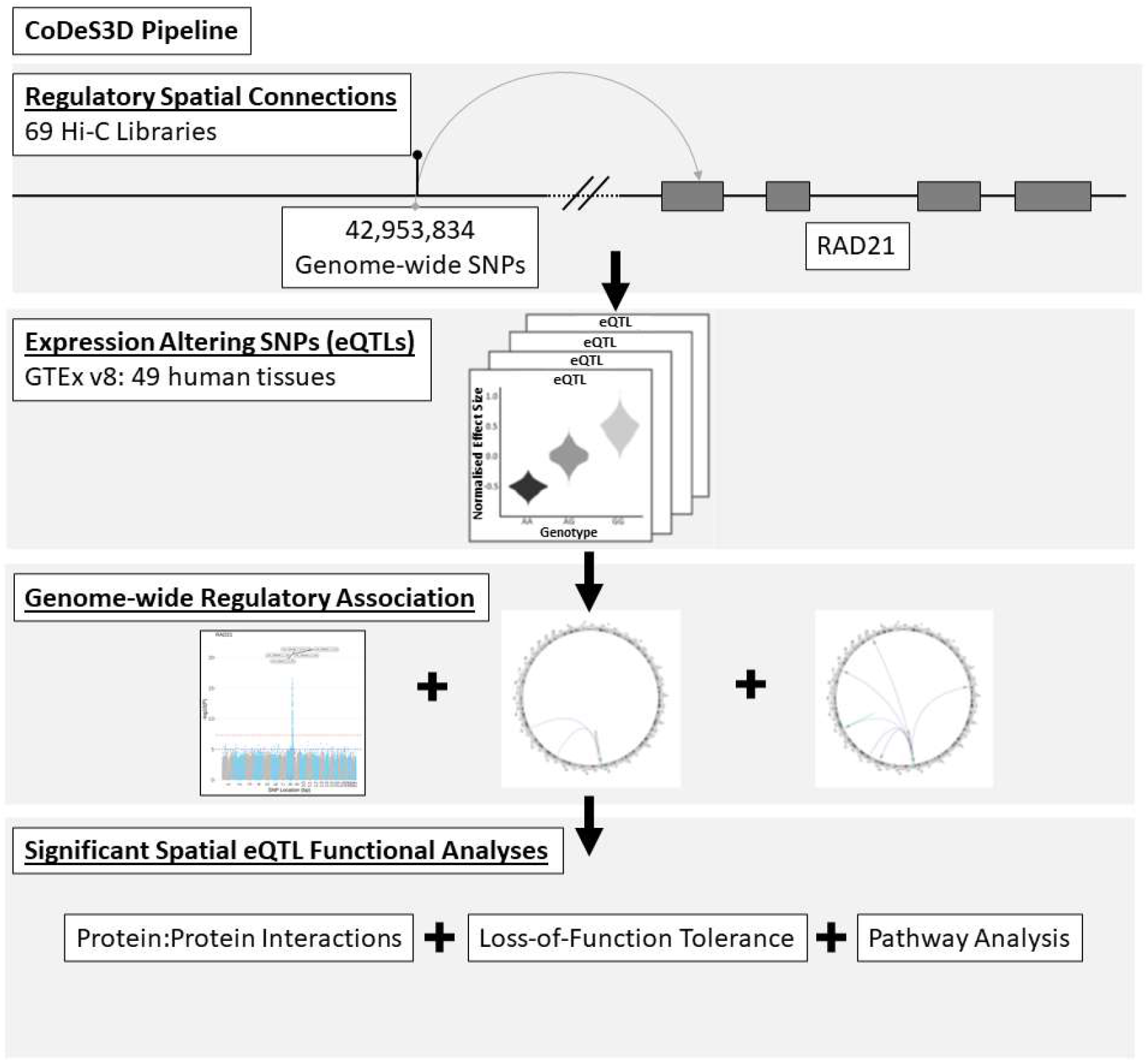
Schematic Representation of the Research Method.

### Identification of SNP-Gene Spatial Relationships

For all variants, spatial regulatory connections were identified through association with transcript levels of *RAD21* (expression Quantitative Trail Locus [eQTL]; GTEx v8^20^) and a confirmed spatial interaction (Hi-C data) using the CoDeS3D algorithm (https://github.com/Genome3d/codes3d-v1)^22,47^. Spatial-eQTL association p-values were adjusted using the Benjamini–Hochberg procedure, and associations with adjusted p-values < 0.05 were deemed spatial eQTL-eGene pairs. Variants with a minor allele frequency below 5% were filtered out due to sample size restrictions within GTEx.

To identify SNP locations in the Hi-C data, reference libraries of all possible Hi-C fragment locations were identified through digital digestion of the hg38 human reference genome with the same restriction enzyme employed in preparing the Hi-C libraries (*i.e*. MboI, HindIII). Digestion files contained all possible fragments, from which a SNP library was created, containing all genome fragments containing a SNP. Next, all SNP-containing fragments were queried against the Hi-C databases to find distal fragments of DNA which spatially connect to the SNP-fragment. If the distal fragment contained the coding region of *RAD21* (GENCODE transcript model GrCH38 positions), a SNP-*RAD21* spatial connection was confirmed. There was no binning or padding around restriction fragments to obtain gene overlap.

Spatial connections were identified from previously generated Hi-C libraries of various origins: 1) Cell lines GM12878, HMEC, HUVEC, IMR90, K562, KBM7, HELA, NHEK, and hESC (GEO accession numbers GSE63525, GSE43070, and GSE35156); 2) tissue-specific data from ENCODE sourced from the adrenal gland, bladder, dorsolateral prefrontal cortex, hippocampus, lung, ovary, pancreas, psoas muscle, right ventricle, small bowel, and spleen (GSE87112); and 3) Tissues of neural origin from the Cortical and Germinal plate neurons (GSE77565), Cerebellar astrocytes, Brain vascular pericytes, Brain microvascular endothelial cells, SK-N-MC, and Spinal cord astrocytes (GSE105194, GSE105513, GSE105544, GSE105914, GSE105957), and Neuronal progenitor cells (GSE52457).

### Identification of regulatory potential of distal SNPs with *RAD21* Spatial Relationships

All SNPs with significant SNP-*RAD21* spatial relationships (FDR < 0.05) were also subsequently tested with the CoDeS3D algorithm^22^ to discover other genes co-regulated by these SNPs.

### Defining mutationally-constrained genes

The human transcriptome consists of genes with varying levels of redundancy and critical function, resulting in some genes being intolerant to loss-of-function (LoF-intolerant) mutation (e.g. *RAD21*). This subset of the human transcriptome are posited to also be more intolerant to regulatory perturbation. The gnomAD catalog^4^ lists 19,704 genes and their likelihood of being intolerant to loss-of-function mutations (pLI), resulting in 3,063 LoF-intolerant, 16,134 LoF-tolerant, and 507 undetermined (15.5% LoF-intolerance, defined as pLI ≥ 0.9)^4^. We tested all genes co-regulated with *RAD21* (as defined above) for LoF-intolerance, comparing our *cis* and *trans*-acting eQTL gene lists for enrichment for LoF-intolerance. This analysis highlights the significance of long-distance gene regulation on otherwise mutationally-constrained (LoF-intolerant) genes, such as *RAD21*.

### Gene Ontology (GO), Pathway Analysis, and Functional Prediction

All genes were then annotated for significant biological and functional enrichment using g:Profiler^48^, which includes the Kyoto Encyclopedia of Genes and Genomes (KEGG) Pathway Database (https://www.kegg.jp/kegg/pathway.html) for pathways and TRANSFAC for transcription factor binding enrichment.

### Protein-Protein Interaction (PPI) Networks

To test for previously known protein-protein interactions associated with our gene lists, all genes were used as inputs into STRING v11 with connection stringency set to low confidence (0.150) and up to 5 interactors added to the network^25^.

## Author Contributions

WS planned the study, performed analyses. WS drafted the manuscript. JAH and JOS acquired the financial support for the project leading to this publication. All authors revised the manuscript and approved the final version.

## Data Availability Statement

The CoDeS3D pipeline and all data used in this paper are available to the public.

**CoDeS3D pipeline**: https://github.com/Genome3d/codes3d-v1

**GTEx portal**: https://www.gtexportal.org/home/

**gnomAD**: https://gnomad.broadinstitute.org/

## Tables

Table 1. A genome-wide search of all 42,953,834 SNPs in dbSNP151 revealed 123 SNPs with significant (FDR < 0.05) regulation of *RAD21* transcript levels. These results were attained through spatial regulatory connections (Hi-C libraries of various tissue origins) confirmed by expression Quantitative Trail Locus (eQTL) analysis (in each tissue available in the GTEx database).

Table 2. All SNPs with significant SNP-*RAD21* spatial-eQTL relationships tested for other genes they co-regulate.

Table 3. All genes were annotated for significant biological and functional enrichment using g:Profiler, which includes the Kyoto Encyclopedia of Genes and Genomes (KEGG) Pathway Database for pathways and TRANSFAC for transcription factor binding enrichment.

Table 4. All genes which comprise the mitotic cohesin and support its loading and unloading, along with three of the seven RAD21-co-regulated genes are LoF-intolerant (pLI < 0.9). We also report the 90% upper bound of the loss-of-function confidence interval (LOEUF).

## References

1. Fudenberg, G. et al. Formation of Chromosomal Domains by Loop Extrusion. Cell Rep. 15, 2038–2049 (2016).

2. Haarhuis, J. H. I. et al. The Cohesin Release Factor WAPL Restricts Chromatin Loop Extension. Cell 169, 693–707.e14 (2017).

3. Horsfield, J. A., Print, C. G. & Mönnich, M. Diverse developmental disorders fromthe one ring: Distinct molecular pathways underlie the cohesinopathies. Front. Genet. 3, 171 (2012).

4. Karczewski, K. J. et al. The mutational constraint spectrum quantified from variation in 141,456 humans. Nature 581, 434–443 (2020).

5. Meier, M. et al. Cohesin facilitates zygotic genome activation in zebrafish. Development 145, dev156521 (2018).

6. Paulsen, J. et al. Long-range interactions between topologically associating domains shape the four-dimensional genome during differentiation. Nat. Genet. 51, 835–843 (2019).

7. Yang, M. et al. Proteogenomics and Hi-C reveal transcriptional dysregulation in high hyperdiploid childhood acute lymphoblastic leukemia. Nat. Commun. 10, 1519 (2019).

8. Achinger-Kawecka, J. et al. Epigenetic reprogramming at estrogen-receptor binding sites alters 3D chromatin landscape in endocrine-resistant breast cancer. Nat. Commun. 11, 320 (2020).

9. Thota, S. et al. Genetic alterations of the cohesin complex genes in myeloid malignancies. Blood 124, 1790–1798 (2014).

10. Rhodes, J. M., McEwan, M. & Horsfield, J. A. Gene regulation by cohesin in cancer: Is the ring an unexpected party to proliferation? Molecular Cancer Research vol. 9 1587–1607 (2011).

11. Cuartero, S., Innes, A. J. & Merkenschlager, M. Towards a Better Understanding of Cohesin Mutations in AML. Frontiers in Oncology vol. 9 867 (2019).

12. Viny, A. D. & Levine, R. L. Cohesin mutations in myeloid malignancies made simple. Current Opinion in Hematology vol. 25 61–66 (2018).

13. Leeke, B., Marsman, J., O’Sullivan, J. M. & Horsfield, J. A. Cohesin mutations in myeloid malignancies: Underlying mechanisms. Experimental Hematology and Oncology vol. 3 13 (2014).

14. Sarogni, P., Pallotta, M. M. & Musio, A. Cornelia de Lange syndrome: From molecular diagnosis to therapeutic approach. Journal of Medical Genetics (2019) doi:10.1136/jmedgenet-2019-106277.

15. Deardorff, M. A. et al. RAD21 mutations cause a human cohesinopathy. Am. J. Hum. Genet. 90, 1014–1027 (2012).

16. Yamamoto, G. et al. Correlation of invasion and metastasis of cancer cells, and expression of the RAD21 gene in oral squamous cell carcinoma. Virchows Arch. 448, 435–441 (2006).

17. Zhou, B. & Guo, R. Genomic and regulatory characteristics of significant transcription factors in colorectal cancer metastasis. Sci. Rep. 8, 17836 (2018).

18. Ahn, J. H., Kim, T. J., Lee, J. H. & Choi, J. H. Mutant p53 stimulates cell invasion through an interaction with Rad21 in human ovarian cancer cells. Sci. Rep. 7, 1–11 (2017).

19. Welter, D. et al. The NHGRI GWAS Catalog, a curated resource of SNP-trait associations. Nucleic Acids Res. 42, D1001–1006 (2014).

20. Aguet, F. et al. The GTEx Consortium atlas of genetic regulatory effects across human tissues. Science (80-.). 369, 1318–1330 (2020).

21. Schierding, W., Horsfield, J. A. & O’Sullivan, J. M. Low tolerance for transcriptional variation at cohesin genes is accompanied by functional links to disease-relevant pathways. J. Med. Genet. 0, jmedgenet-2020-107095 (2020).

22. Fadason, T., Schierding, W., Lumley, T. & O’Sullivan, J. M. Chromatin interactions and expression quantitative trait loci reveal genetic drivers of multimorbidities. Nat. Commun. 9, 5198 (2018).

23. Xie, J., Yin, H., Nichols, T. D., Yoder, J. A. & Horowitz, J. M. Sp2 is a maternally inherited transcription factor required for embryonic development. J. Biol. Chem. 285, 4153–4164 (2010).

24. Chin, C. V. et al. Cohesin mutations are synthetic lethal with stimulation of WNT signaling. Elife 9, 1–21 (2020).

25. Szklarczyk, D. et al. STRING v11: Protein-protein association networks with increased coverage, supporting functional discovery in genome-wide experimental datasets. Nucleic Acids Res. 47, D607–D613 (2019).

26. Sowa, M. E., Bennett, E. J., Gygi, S. P. & Harper, J. W. Defining the Human Deubiquitinating Enzyme Interaction Landscape. Cell 138, 389–403 (2009).

27. Völkel, S. et al. Zinc Finger Independent Genome-Wide Binding of Sp2 Potentiates Recruitment of Histone-Fold Protein Nf-y Distinguishing It from Sp1 and Sp3. PLOS Genet. 11, e1005102 (2015).

28. Yeh, C. et al. The Deubiquitinase USP37 Regulates Chromosome Cohesion and Mitotic Progression. Curr. Biol. 25, 2290–2299 (2015).

29. He, X. et al. USP13 interacts with cohesin and regulates its ubiquitination in human cells. J. Biol. Chem. 296, 100194 (2021).

30. Dou, N. et al. USP32 promotes tumorigenesis and chemoresistance in gastric carcinoma via upregulation of SMAD2. Int. J. Biol. Sci. 16, 1648–1657 (2020).

31. Guo, Y. et al. Comprehensive ex vivo transposon mutagenesis identifies genes that promote growth factor independence and leukemogenesis. Cancer Res. 76, 773–786 (2016).

32. Fang, C., Rao, S., Crispino, J. D. & Ntziachristos, P. Determinants and role of chromatin organization in acute leukemia. Leukemia vol. 34 2561–2575 (2020).

33. Paulding, C. A., Ruvolo, M. & Haber, D. A. The Tre2 (USP6) oncogene is a hominoid-specific gene. Proc. Natl. Acad. Sci. U. S. A. 100, 2507–2511 (2003).

34. Shen, C. et al. Calcium/calmodulin regulates ubiquitination of the ubiquitin-specific protease TRE17/USP6. J. Biol. Chem. 280, 35967–35973 (2005).

35. Panigrahi, A. K., Zhang, N., Mao, Q. & Pati, D. Calpain-1 Cleaves Rad21 To Promote Sister Chromatid Separation. Mol. Cell. Biol. 31, 4335–4347 (2011).

36. Getz, G. et al. Integrated genomic characterization of endometrial carcinoma. Nature 497, 67–73 (2013).

37. Ley, T. J. et al. Genomic and epigenomic landscapes of adult de novo acute myeloid leukemia. N. Engl. J. Med. 368, 2059–2074 (2013).

38. Hu, W. et al. Downregulation of USP32 inhibits cell proliferation-migration and invasion in human small cell lung cancer. Cell Prolif. 50, 50 (2017).

39. Akhavantabasi, S. et al. USP32 is an active, membrane-bound ubiquitin protease overexpressed in breast cancers. Mamm. Genome 21, 388–397 (2010).

40. Kjällquist, U. et al. Exome sequencing of primary breast cancers with paired metastatic lesions reveals metastasis-enriched mutations in the A-kinase anchoring protein family (AKAPs). BMC Cancer 18, 174 (2018).

41. Zaharija, B., Samardžija, B. & Bradshaw, N. J. The TRIOBP Isoforms and Their Distinct Roles in Actin Stabilization, Deafness, Mental Illness, and Cancer. Molecules (Basel, Switzerland) vol. 25 (2020).

42. Xu, S. et al. RCN1 suppresses ER stress-induced apoptosis via calcium homeostasis and PERKCHOP signaling. Oncogenesis 6, e304–e304 (2017).

43. Bonne, A. et al. Mapping of constitutional translocation breakpoints in renal cell cancer patients: identification of KCNIP4 as a candidate gene. Cancer Genet. Cytogenet. 179, 11–18 (2007).

44. Brenner, D. R. et al. Identification of lung cancer histology-specific variants applying Bayesian framework variant prioritization approaches within the TRICL and ILCCO consortia. Carcinogenesis 36, 1314–1326 (2015).

45. Phan, D. et al. Identification of Sp2 as a Transcriptional Repressor of Carcinoembryonic Antigen-Related Cell Adhesion Molecule 1 in Tumorigenesis. Cancer Res. 64, 3072–3078 (2004).

46. Zhu, Y. et al. Sp2 promotes invasion and metastasis of hepatocellular carcinoma by targeting TRIB3 protein. Cancer Med. 9, 3592–3603 (2020).

47. Schierding, W. et al. GWAS on prolonged gestation (post-term birth): analysis of successive Finnish birth cohorts. J. Med. Genet. 55, 55–63 (2018).

48. Raudvere, U. et al. g:Profiler: a web server for functional enrichment analysis and conversions of gene lists (2019 update). Nucleic Acids Res. 47, W191–W198 (2019).

